# Exploring the embodied mind: functional connectome fingerprinting of meditation expertise

**DOI:** 10.1101/2023.12.06.570128

**Authors:** Sébastien Czajko, Jelle Zorn, Loïc Daumail, Gael Chetelat, Daniel Margulies, Antoine Lutz

## Abstract

**BACKGROUND:** Short mindfulness-based interventions have gained traction in research due to their positive impact on well-being, cognition, and clinical symptoms across various settings. However, these short-term trainings are viewed as preliminary steps within a more extensive transformative path, presumably leading to long-lasting trait changes. Despite this, little is still known about the brain correlates of these meditation traits.

**METHOD:** To address this gap, we investigated the neural correlates of meditation expertise in long-term Buddhist practitioners, comparing the large-scale brain functional connectivity of 28 expert meditators with 47 matched novices. Our hypothesis posited that meditation expertise would be associated with specific and enduring patterns of functional connectivity present during both meditative (open monitoring/open presence and loving-kindness compassion meditations) and non-meditative resting states, as measured by connectivity gradients.

**RESULTS:** Applying a support vector classifier to states not included in training, we successfully decoded expertise as a trait, demonstrating its non-state-dependent nature. The signature of expertise was further characterized by an increased integration of large-scale brain networks, including the dorsal and ventral attention, limbic, frontoparietal and somatomotor networks. The latter correlated with a higher ability to create psychological distance from thoughts and emotions.

**CONCLUSION:** Such heightened integration of bodily maps with affective and attentional networks in meditation experts could point toward a signature of the embodied cognition cultivated in these contemplative practices.

## INTRODUCTION

Short 8-week mindfulness-based interventions (MBIs), which are routinely used in various clinical and educational settings, can positively impact well-being and cognition (1), and decrease clinical symptoms, in particular in mood disorders (2,3, for a review see 4). MBI can induce functional changes in the neural processes underlying affect and attention (5,6) which are not always associated with structural changes (7), the latter being reported following several years of meditation training (8). According to traditional meditation theories, these short-term training effects are only preliminary within a more transformative path leading to long-lasting trait changes in cognition and self-related processes (9). We previously reported that a sample of long-term Tibetan Buddhist practitioners had better pain regulation capacity (10), lower trait measures of depression, anxiety, pain catastrophizing, and reported higher cognitive defusion skill (11), different pain regulatory strategies (12) and pro-social dispositions (13) than a group of matched novices. Despite the potential therapeutic and scientific value of long-term meditation expertise, little is still known about its neurophysiological mechanisms and conflicting findings exist regarding the preliminary research (14,15). The purpose of the present study is to investigate the neural correlates of meditation expertise in this sample of long-term Tibetan Buddhist meditators and novices, as measured by changes in the organization of intrinsic connectivity networks in the brain.

The mental training leading to this expertise can be conceptualized, in this tradition, as the process of getting familiarized with one’s mind by practicing various meditative techniques. The developmental trajectory starts typically by cultivating mindfulness and compassion practices for several years (see SM for details) before gradually transitioning into non-dual meditation such as Open Presence (OP) (10). This practice aims at exploring and gaining insights into the constructive and transient nature of basic cognitive structures such as time, self, and subject–object orientation. OP is said to induce a minimal phenomenal state of consciousness where the intentional structure involving the duality between object and subject is attenuated, as captured by the notion of non-duality (16,17). OP meditation is said to have long-term impact on cognition and perception as a trait such as the formal non-dual meditation is gradually and spontaneously integrated in daily life (18). From this description of this meditation expertise, our overall hypothesis was that experts’ resting state (RS) brain activity would resemble their activity during meditative states (here open monitoring (OM) and loving-kindness and compassion (LKC)), and particularly of the non-dual ones (i.e. OP). We predict that any long-lasting, trait-like changes in experts will be associated with specific changes compared to novices in large-scale brain functions detectable both during meditative states and at rest during non-meditative states.

To investigate our hypothesis of a trait-like effect of long-term meditation practice on the intrinsic functional organization of the cerebral cortex (19), we employed diffusion embedding, also known as connectivity gradients (20,21). This method appears to be promising to tackle our question for several reasons. Firstly, its data-driven approach overcomes the limitations associated with the hypothesis-driven approaches traditionally used in the field. Secondly, it can detect connectivity differences missed by Independent Component Analysis (22). This technique can reveal multiple dimensions of cortical organization, with the first dimension describing the cognitive hierarchy (19,20), starting from sensory cortices and ending with transmodal regions such as the default-mode network (DMN). The second gradient separates visual regions from the other networks (20), and the third gradient spans the multiple-demand network and the networks at opposite end (23). Previous studies have demonstrated that these gradients can be influenced by various factors, including disorders such as depression (24) and autistic spectrum disorder (25), as well as cognitive and psycho-affective training (26).

We employed these three gradients as continuous coordinates in a 3D space, and computed the eccentricity of all brain vertices located in this 3D space as their distance to the barycenter (22,26). Building on this approach, we used a set of specialized measures that quantify the dispersion within and between functional communities in a connectivity-derived manifold (22). As such, they were chosen as our best candidates to characterize changes associated with meditation expertise in the brain connectome as a whole (red in Figure 1C), and within and between these functional communities (green and black in Figure 1C), (22,26, see Methods). Given the paucity of data in the literature on meditation expertise and novelty of the gradient connectivity method, further developing a more refined functional hypothesis is challenging. However, one can identify brain networks candidates which may be impacted within the connectome. Meditation has been associated with brain structural and functional changes mainly in frontal and limbic networks (27,28) with the insula and anterior cingulate cortex, part of the salience network (SN), being the regions most sensitive to meditation training according to a meta-analysis (29). Studies on meditation traits and functional connectivity (30,31) or differences between long-term meditation practitioners and novices (14,15,32) have also shown that individuals with meditation experience exhibit reduced connectivity between the DMN and frontoparietal network (FPN)/SN. Yet, these effects remain inconsistent across studies, as meditation training has also been linked to increased connectivity between the DMN and FPN/SN (33,34). A recent study reported effects of various forms of meditation training on the connectome: training in attentional family meditation akin to FA/OM increased functional segregation of regions, including parietal and posterior insular regions, indicating that these networks are functionally different from the rest of the cortex (26). Conversely, training in perspective involving meta-cognitive and perspective taking on self and others resulted in increased functional integration of these regions with other brain networks (26). As both trainings potentially share some features with OP (e.g. meta-awareness and dereification), it is difficult, based on Valk et al.’s study alone, to predict whether the eccentricity of experts would decrease or increase. Moreover, contrary to these two trainings, OP aims at suspending the subject-object duality, an effect that is a common outcome of psychedelics inducing ego dissolution (9,35). As the latter effect has been shown to decrease the segregation of large-scale networks within the first gradient (36,37), it is plausible that experts would exhibit lower eccentricity compared to novices.. In our exploratory study, we employed a machine learning approach to uncover the hidden patterns of brain connectivity characterizing meditation expertise: we trained a support vector classifier (SVC) on a subset of a given state to distinguish experts from novices, and then tested its ability to both decode the same state and to generalize to the other states. The prediction from our hypothesis was that if the expertise effect was an enduring dynamical characteristic present in every state, the SVC should be able to generalize to the other states as well. We then used these measures to characterize expertise in multivariate analyses (Figure 2A, 2E), followed by exploratory univariate analyses (Figure 2B). We further examined whether these measures could also predict psychometric and meditative trait measures.

**Figure 1.**
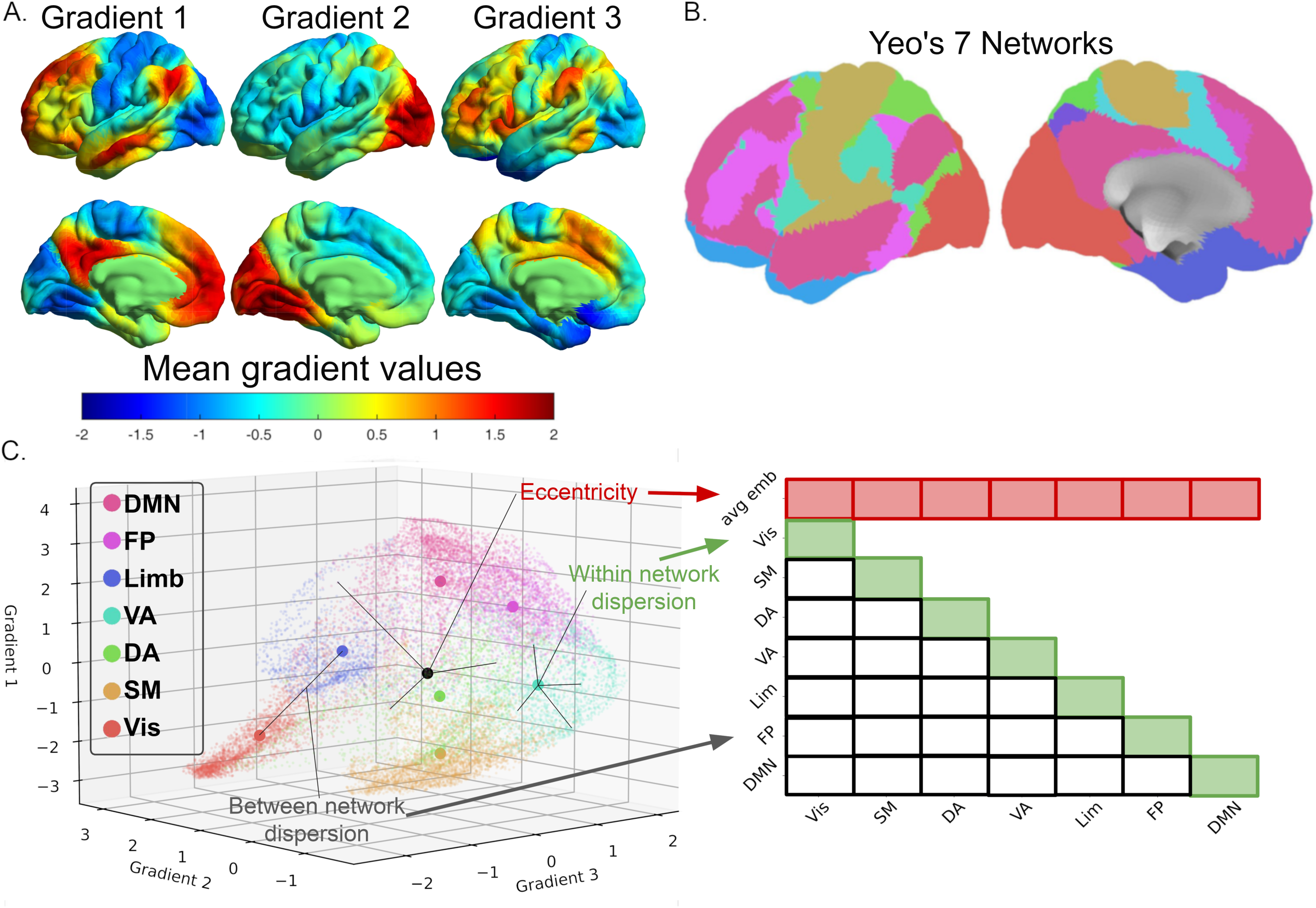
Gradients and eccentricity maps (A) The first gradient (left) denotes the cognitive hierarchy of the brain, ranging from the unimodal cortex to the DMN. The second gradient (middle) differentiates the visual cortex from other networks, while the third gradient (right) segregates the limbic network. (B) Visualization of the cortical parcellation (63) used in (Figure 1C). The color code corresponds to the one in Figure 1C (C) Visualization of dispersion metrics within the three-gradient space (right). The eccentricity value of a vertex corresponds to the Euclidean distance from the barycenter (depicted by a black dot) of the 3D space. For each individual 3D map, we computed an eccentricity map defined by the Euclidean distance from each vertex to the individual barycenter of the 3D space (21,37). These maps reflect the integration (low eccentricity) and segregation (high eccentricity) within the connectome for each vertex of each participant. We then quantified the dispersion metrics, which represents the segregation of large-scale networks (21). The within-network dispersion is calculated as the sum squared Euclidean distance of network vertices to the network barycenter. Between-networks dispersion is quantified as the Euclidean distance between network barycenters (21). Visual explanation of the dispersion metrics (left). The first row of the matrix shows the average vertex-wise eccentricity of each network, referred to as the average embedding (21). The remaining dispersion metrics display the within-network dispersion (diagonal in green) and the between-network dispersion (the remaining black squares). A set of dispersion measures was computed for every participant and every state, and used to decode meditation expertise using a Support Vector Classifier (SVC) algorithm.

**Figure 2.**
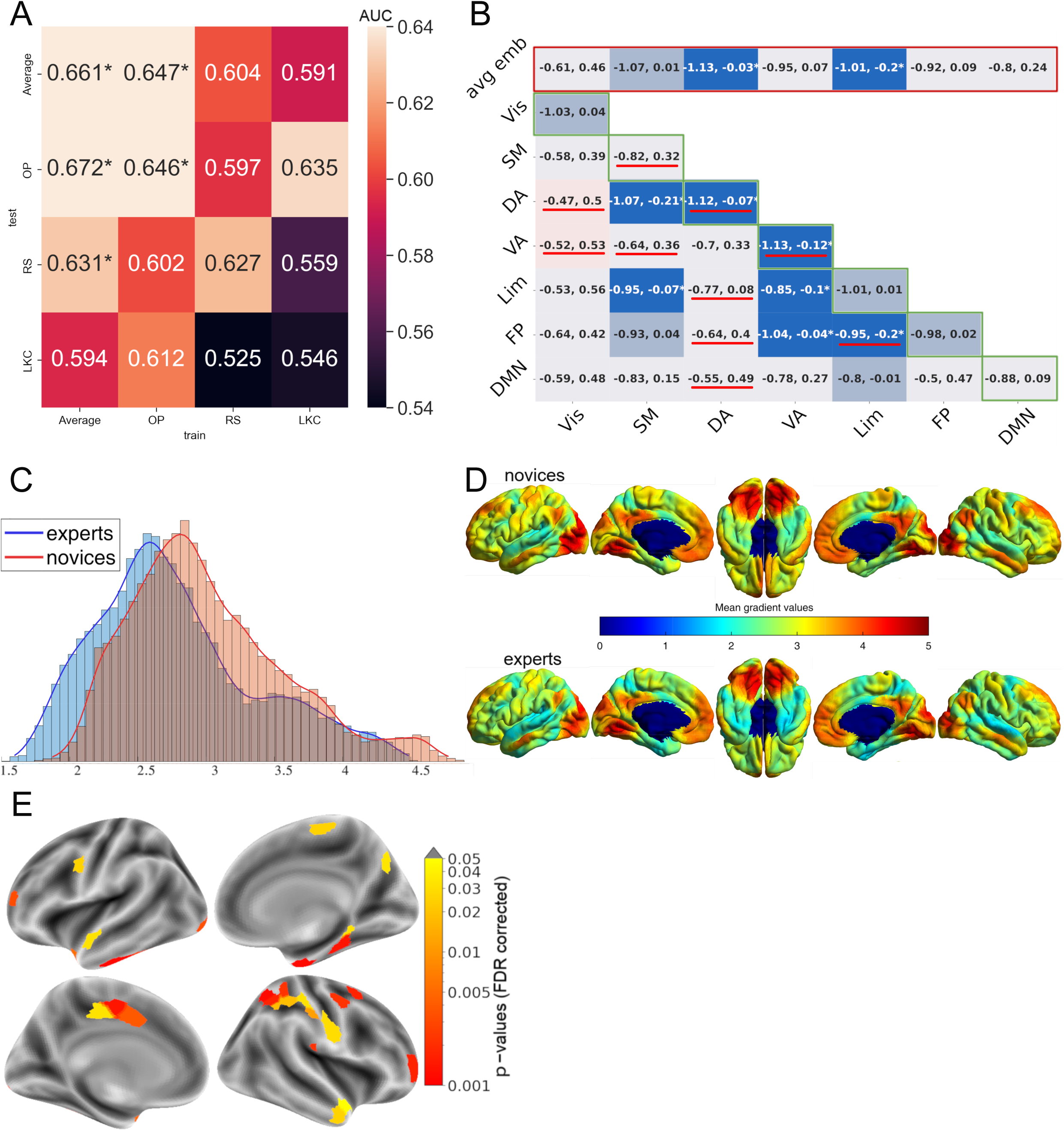
Effects of meditation expertise on eccentricity maps and dispersion metrics (A) A stochastic gradient descent classifier was trained on the dispersion metrics (Figure 2B, Figure 1C) to decode expertise. The training was performed on a subsample of the three states or the average of the three states. The classifier was tested on the same metrics of the remaining sample, either from the same state or from a different state, using AUC. The SVC could decode expertise using the dispersion metrics of the average state, and the OP state, but not in LKC and RS. The decoder trained on the average state could predict the expertise from resting state and OP data. A star displays a significant effect (p<0.05). (B) Comparison of the dispersion metrics of the averaged state between experts and novices. Significance was tested using bootstrap-t tests (39), and 95% confidence intervals are plotted in each box. The color blue (respectively red) indicates that the average eccentricity value was lower (respectively higher) for experts than novices. Bright-colored boxes indicate significant tests (p<0.05), medium-colored boxes indicate a trend (0.1>p>0.05), while light-colored boxes indicate non-significant tests (p>0.1). The tests here are exploratory and thus are not corrected for multiple comparisons. The results indicate that several metrics were lower for experts compared to novices. Here, a higher between-network dispersion reflects a weaker connectivity between two networks and a higher within network dispersion, a lower connectivity between the vertices of a given network. The red underlines indicate which dispersion metrics contributed significantly to the decoding of expertise (Figure 2A). (C) Histograms of the averaged state eccentricity values for experts and novices. There was only a trend for the mean eccentricity to be lower for experts than novices (t_boot_=0.44; p=0.09). (D) The eccentricity map describes a continuous coordinate system, where a lower value signifies that the vertex is closer to the barycenter of the 3D space. For both groups, sensory regions, including the visual and somatomotor cortex, and the DMN were the least integrated regions. Although the average maps of experts and novices are similar overall, the eccentricity values tended to be visually lower for experts than novices. (E) Surface-wide statistical comparisons between novices and experts after averaging RS, OP and LKC eccentricity maps. The voxels within a parcel were used to decode expertise to get an AUC value for each parcel. This analysis characterizes expertise at a finer scale than large-scale networks. Significant parcels belong mainly to the sensorimotor (SM), dorsal attention (DA), ventral attention (VA), and limbic (Lim) networks. (F) This violin plot displays the averaged eccentricity of the significant parcels in Figure 2E, which was lower for experts than for novices (t(68)=-2.54; p=0.014).

Investigating expert meditators could thus help develop hypotheses or theoretical understanding about the long-term developmental trajectory of meditation and, possibly, non-dual states.

## METHODS

### Participants

Participants were recruited for a cross-sectional study on mindfulness effects, part of the Brain and Mindfulness project in Lyon (2015–2018). Participants included novices and long-term meditation practitioners (referred to as ‘experts’), who were recruited through multiple screening stages (for details, see the Brain & Mindfulness Project Manual (38)). More precisely, meditation practitioners should have accumulated ≥ 10000 hours of experience, followed at least one formal 3-year meditation retreat, and have a regular daily practice of ≥ 45 minutes in the year preceding the study. 75 cognitively normal participants aged 35-66 (SD 7.7) including 28 expert meditators and 47 control participants (referred to as ‘novices’) matched on age and gender (p>0.5, see Table 1) were included in this study. Novices attended a one-weekend meditation training program prior to any measurement to get familiarized with the meditation techniques. Inclusion and exclusion criteria have been previously reported (38) (see SM). Finally, subjects had to be affiliated to a healthcare system. All participants received information on the experimental procedures during a screening session, and provided informed written consent. The study and its analyses were approved by the regional ethics committee on Human Research (CPP Sud-Est IV, 2015-A01472-47). The sample size of this study was powered for a fMRI pain paradigm presented in another publication (10). While the power analysis for the fMRI data was based on a group size of 25 participants (plus 3 participants to accommodate for artifactual data), we oversampled the novice group to increase our power for correlation analyses with questionnaire measures. After excluding participants who exhibited more than 0.3mm/degree movement to control for potential motion effects (2 experts and 3 novices), the analysis was conducted with a reduced sample size of 70 participants (39).

**Table 1:**
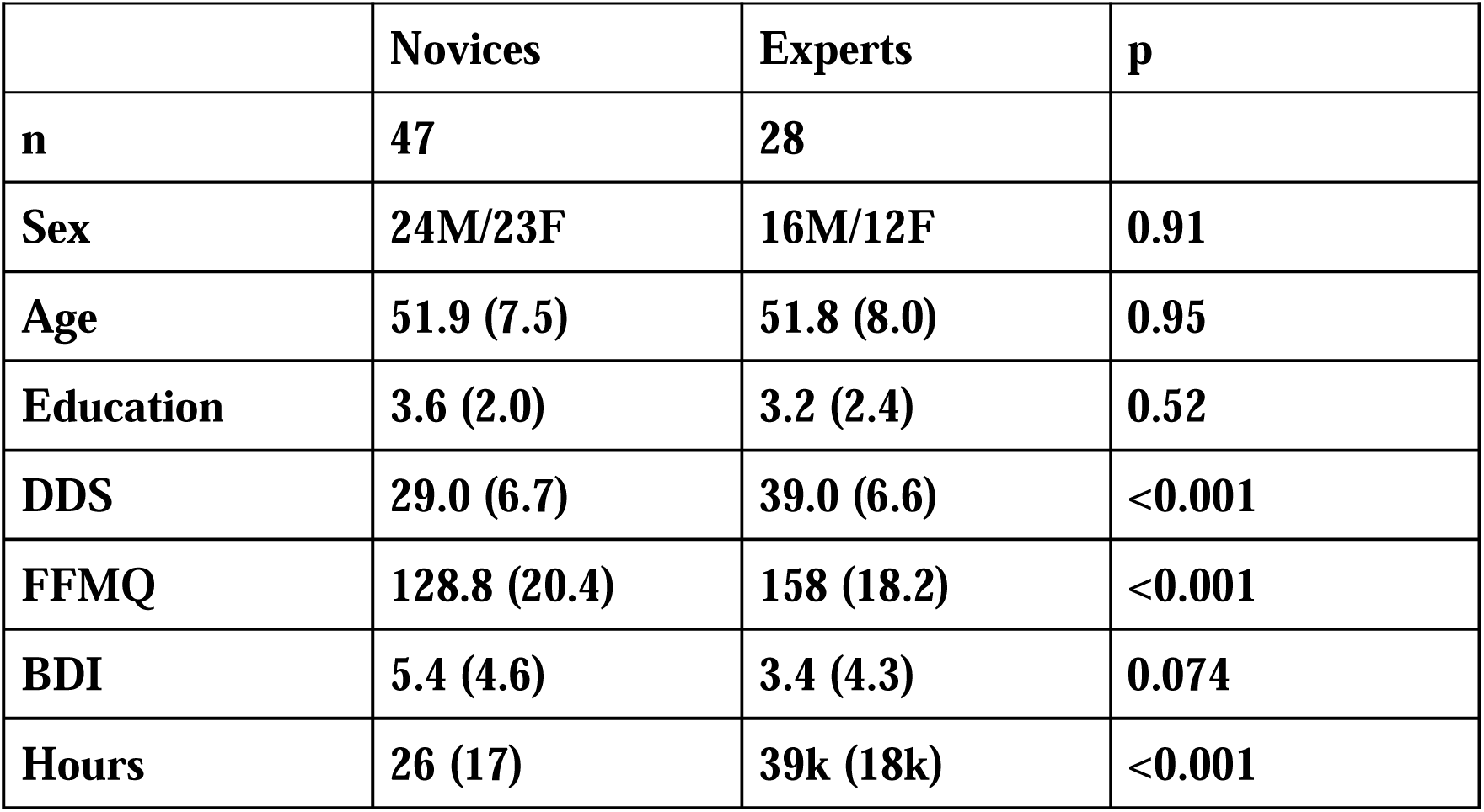
Demographics comparison between experts and novices. Continuous variables are presented as mean (standard deviation) and their p-values were calculated using t-test. For sex, the p-value was calculated using a chi-squared test. DDS: Drexel Defusion Scale. FFMQ: Five Facets Mindfulness Questionnaire. BDI: Beck Depression Inventory.

### Paradigm

All participants attended a single fMRI session in which we first acquired their structural image. We then acquired functional scans, starting with a RS. We also acquired meditative states of LKC meditation. In addition, for novices, we acquired states of OM meditation, and for experts we acquired states of OP meditation (see SM for a description of the meditation practices). All states lasted 10 minutes. The order of acquisition of the two meditative states was random. For the present study, we used three psychometric scales, which are Drexel defusion scale (DDS) (40), Five facets mindfulness questionnaire (FFMQ) (41), and Beck depression inventory (BDI) (42) (for details, see SM).

### Data acquisition and preprocessing

Data was collected on a 3T Siemens Prisma scanner. Functional data was acquired with EPI (TR=2100ms, TE=30ms, 39 slices, voxel size 2.8×2.8×3.1mm3). Structural scans were T1w (1mm iso), T2w (1mm iso) and T2*w (1mm iso). Preprocessing used fMRIprep v1.2.6 (43). This included motion correction, co-registration, normalization to MNI space, CompCor for physiological noise removal, ICA-AROMA denoising, and FreeSurfer surface reconstruction (see SM for details).

### Connectome gradient construction

The construction of the functional connectome gradient followed the procedures detailed in Hong et al. (2019) (25) and in the SM.

### 3D gradient metrics

To investigate multidimensional differences in cortical organization, we focused on the first three components, which explained over 50% of total variance. We combined these gradients by forming a 3D space (22,26), where each gradient constitutes an axis of this space described in Figure 1C. From there, we computed the eccentricity representing the level of integration for any given vertex; the most integrated vertices have the lowest eccentricity values (dorsal attention (DA), ventral attention (VA)), while highly specialized (DMN) or unimodal vertices exhibit low integration relative to the rest of the brain and therefore also have low eccentricity values. Additionally, to compare our findings with more conventional connectivity approaches (44), and suggested by recent studies (22,26), we derived two other metrics, the between-network dispersion and the within-network dispersion. A higher dispersion of these metrics indicates respectively a lower connectivity between networks or within that network.

### Statistical analyses

To identify the state that best characterizes expertise according to our hypothesis, we trained classifiers on the dispersion metrics matrix to predict expertise using scikit-learn (45) with a modified-huber loss and 5-fold cross-validation repeated 5000 times. We used the area under the curve score rather than accuracy to avoid bias from unbalanced samples. Once we had identified this state, we conducted exploratory post-hoc tests using Studentized bootstrap-t tests with 10.000 repetitions (46) to characterize which dispersion metrics were different between experts and novices. Then, to characterize these differences at a finer scale, we compared eccentricity values between experts and controls on the surface using a ridge classifier with a 3-fold cross-validation scheme (45) repeated 300 times on each of the 400 parcels of the Schaeffer Atlas (47). Significant parcels were identified using the same methodology that was applied in the searchlight classification informative region mixture model (48). Subsequent p-values were adjusted using False Discovery Rate correction (49). Surface-based linear models computed with SurfStat (http://www. math.mcgill.ca/keith/surfstat/) and corrected for family-wise errors using random field theory (pFWE < 0.05) are available in SM (Figure S1). Finally, to disentangle collinear demographic factors, we used a back-to-back regression (B2B) (50), finding DDS score had a significant contribution to dispersion measures (details in SM).

## RESULTS

### DECODING ANALYSIS

We hypothesized that the large-scale fMRI connectomics measures would be modulated by trait-like effects of expertise not only during meditative states but also at rest during non-meditative states. To test this hypothesis, we used a machine learning approach. We trained a support vector classifier (SVC) to distinguish experts from novices using the dispersion metrics described in Figure 1C on a subset of participants in a given state, and then tested for its ability to decode both the same state and to generalize the decoding to the other states. Our rationale was that if the expertise effect was an enduring dynamical characteristic present in every state, the SVC should be able to generalize to the other states as well. Specifically, the state with the least amount of noise around the effect of expertise should be the most susceptible to generalization when tested on another state (Figure 2A). We were able to decode expertise only for the OP state (AUC=0.646; p=0.027), but we were not able to generalize its classification on the other states. Next, to better measure the effect of expertise, we averaged all three states together, and again trained the classifier using the average state’s dispersion metrics. As expected, the model demonstrated significant expertise decoding ability when trained on the average state and subsequently tested on the remaining test set (AUC=0.661; p=0.021). Interestingly, unlike when trained on OP, the model was also able to generalize its classification ability when tested on different states, specifically RS (AUC=0.631; p=0.046) and OP (AUC=0.672; p=0.02). In line with our hypothesis, this indicates that the impact of expertise on large-scale networks, as measured by gradient dispersion, is evident across multiple distinct cognitive states and is more accurately represented by their average. Next, we will now focus on describing the specific differences between novices and experts within this averaged state.

### AVERAGED STATE DISPERSION ANALYSIS

To further characterize the averaged state that best captured the fingerprint of meditation expertise, we examined the dispersion metrics of this averaged state using both a 3D space exploratory analysis (Figure 2A) and a surface-based analysis (Figure 2B). Following the exploratory analysis, we found a significant decrease in averaged eccentricity for the experts of the DA using a Studentized bootstrap (t_boot_=0.54; p=0.046), limbic (t_boot_=0.61; p=0.017) networks, a trend toward significance for the somatomotor network (t_boot_=0.51; p=0.058) and within the dorsal (t_boot_=0.57; p=0.029) and VA (t_boot_=0.6; p=0.02) networks. Additionally, we observed a decrease in dispersion between the limbic network and the somatomotor (t_boot_=0.52; p=0.038), VA (t_boot_=0.48; p=0.039), and fronto-parietal (t_boot_=0.56; p=0.021) networks, as well as between the VA and the fronto-parietal (t_boot_=0.51; p=0.037) networks, and between the DA network and the somatomotor cortex (t_boot_=0.62; p=0.012). Regarding the surface-based analysis, we performed a multivariate analysis using the averaged state’s eccentricity on each parcel of the Schaefer 400 atlas (47) to decode expertise (Figure 2E). Significant parcels belonged mainly to the somatomotor and DA of the right hemisphere. In order to quantify this difference of eccentricity between experts and novices, we averaged the eccentricity within all significant parcels and compared the two groups based on this average. This averaged eccentricity was lower for experts than for novices (t(68)=-2.54; p=0.014).

To investigate the behavioral relevance of these group differences, we then studied the individual contribution of various features, including sex, age, group, hours of meditation practice in life, and trait psychometric measures (DDS, BDI, FFMQ) to decode the dispersion metrics. To do so, we fitted a B2B model to control for the co-variance between features while optimizing the linear combination of dispersion metrics to detect the encoding information (50, Figure 3A). The output of this model is a set of beta coefficients, one for each feature. Here, only the DDS, a scale reflecting a person’s capacity to cognitively defuse thoughts, and emotions, yielded a significant contribution to the decoding of meditation expertise (β=0.31; p=0.043). We then applied the same B2B model to each dispersion metric individually, meaning that we used all previous features to predict dispersion metrics. We only present this exploratory analysis for DDS, as it was the only scale to demonstrate a significant relationship (Figure 3B). Our goal was to identify which eccentricity metrics predicted by the set of features exhibited a significant contribution from the DDS. The results indicate that the DDS can predict primarily the same metrics that are significant in Figure 2B. A higher DDS trait score is associated with less dispersion in these metrics. More specifically, a higher DDS score was associated with lower average eccentricity in somatomotor (β=0.13; p=0.023), VA (β=0.09; p=0.046) and limbic (β=0.11; p=0.034) networks. A higher DDS score was also associated with less dispersion between the somatomotor network on one side and the DA (β=0.21; p=0.006), VA (β=0.1; p=0.038), limbic networks (β=0.15; p=0.016), and DMN (β=0.11; p=0.029) on the other side, and between the limbic and the VA networks (β=0.11; p=0.036), and between the FP network and the DMN (β=0.15; p=0.015). However, contrary to Figure 2B, a higher DDS score was not associated with a change of dispersion within networks. To summarize, our analysis suggested that the capacity to put psychological distance between thoughts and emotions was associated with reduced network dispersion between and across specific networks, largely overlapping with the expert-related trait signature (Figure 2B), indicating a potential link between trait-like measures and neural activity during meditation. These findings shed light on the neural mechanisms underlying expertise effects in meditation and highlight the importance of considering state-averaging approaches in future studies.

**Figure 3.**
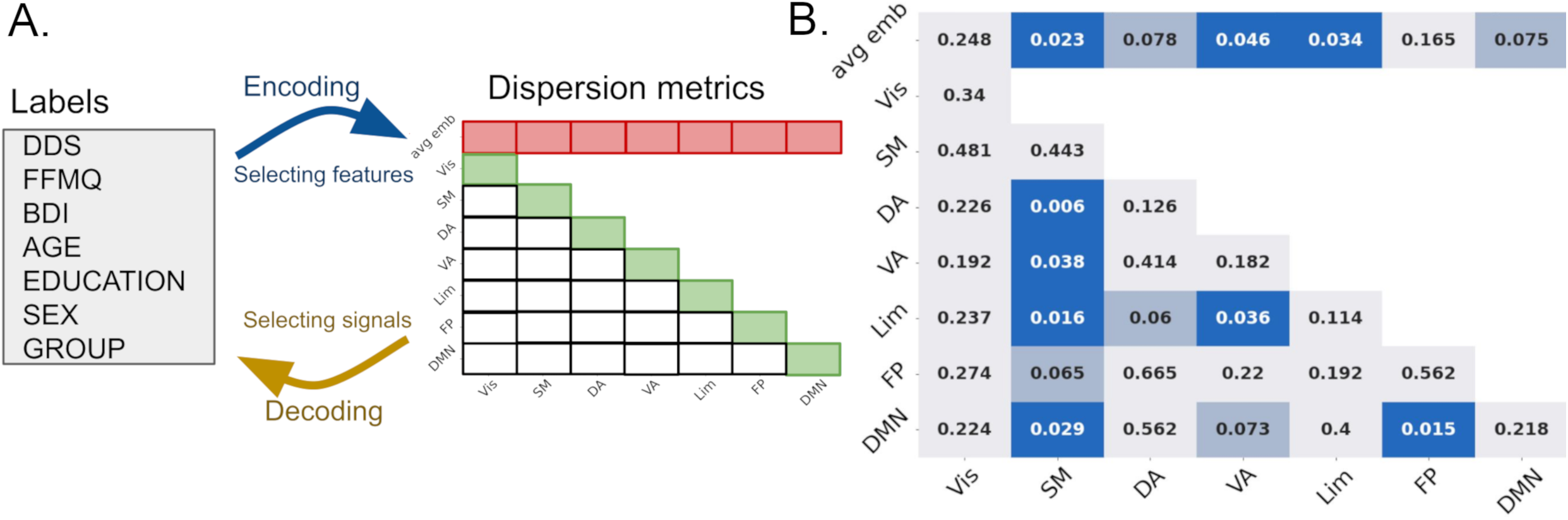
Traits factor analysis (A) The B2B regression method was computed, using the indicated labels as features and dispersion metrics from the average state as signals. An encoder was used on top of a decoder to determine the importance of features despite their shared covariance. The DDS was found to be the only significant feature (β=0.31; p=0.043). (B) During an exploratory analysis, the dispersion metrics were used to predict DDS scores using B2B on each metric. The color code is similar to Figure 2B, with the difference that blue corresponds to a negative correlation and red to a positive correlation between the DDS and the corresponding dispersion metric. For instance, it shows that the more the somatomotor network was integrated to other networks , the higher was the DDS score.

## DISCUSSION

In this study, we explored the neural correlates of meditation expertise as measured by changes in the organization of intrinsic functional connectivity networks in the brain. First, we only managed to decode the groups when it was trained and tested on the OP state, but not on RS or LKC states, suggesting that this state was functionally the most different between groups. Yet, this pattern did not generalize as a trait, meaning that the SVC weights likely captured a trait-by-state effect. Next, we repeated the same procedure on the average of the three states, as the trait effect has been characterized by low-variability functional connectivity (51). If OP-related group differences were reflecting only a state effect, the predictability should decrease, because noise was added during the averaging. If, instead, averaging the states reduced noise by repeating a trait-like feature, then its ability to generalize to other states should increase. We found some evidence for the latter (Figure 2A), suggesting that the average eccentricity was the best characterization of a trait-like effect in our sample.

Subsequent analyses specified further the fingerprint of meditation expertise on the eccentricity map (22,26), which reflects the functional integration (low eccentricity) and segregation (high eccentricity) along a scalar value . Experts exhibited reduced average eccentricity in DA and limbic networks, and, at tendency in the somatomotor cortex, suggesting that these networks were more integrated within the cognitive hierarchy for experts, allowing for enhanced information exchange with other networks (22,26). In line with these findings, our multivariate analysis on the Schaefer atlas parcels revealed clusters, whose eccentricity was lower among the expert group (Figure 2B) in the right parahippocampal gyrus, premotor gyrus, and supplementary motor area. Experts also demonstrated a more reduced within-network dispersion in the DA and VA networks compared to novices, aligning with previous reports on meditation traits (15,30,31), suggesting an enhanced spread of information within these networks, as their vertices exhibit stronger connectivity. Finally, for experts only, the limbic network displayed increased connectivity with the somatomotor, VA, and FP networks, the somatomotor cortex exhibited stronger connectivity with the DA network, while the VA network demonstrated enhanced connectivity with the FP network. This decrease of eccentricity in experts is consistent with the results reported by Valk et al. (26), where perspective training led to similar reductions in eccentricity. However, they contrast with findings from the same study that observed increased eccentricity following attention training, suggesting that different meditation and cognitive training practices may target and modulate distinct neural mechanisms. Interestingly, similar but much more pronounced patterns of global increased integration have been identified in studies investigating the acute effects of psychedelics using diffusion map embedding (36,37). Consistent with our hypothesis, this compression of the cortical hierarchy might be associated with the lessening of self-related/discursive processes during non-dual meditation akin to OP meditation. Similarly, previous studies have observed increased integration in long-term meditation practitioners’ brains using different methodologies, such as graph analysis (52) and diffusion-weighted imaging (53–55). Numerous studies have similarly highlighted the role of these attention and affective brain networks during meditation practices (27,44,56). The functional coupling of these networks with the somatomotor network in meditation is more rarely reported (57), even if it is consistent with the embodied nature of this practice (58,59). This finding pointed toward an important functional modulation of the somatomotor cortex in meditation practice, which have often not been utilized as seeds or networks of interest in previous ICA studies.

We reported that several metrics capturing the meditation expertise fingerprint were correlated with the ability to create psychological distance between thoughts and emotions, as measured by the DDS. These correlations were assessed while considering the co-variation of all metrics included in the demographic table (Table 1), including expertise. Specifically, and in line with the trait fingerprint, a higher DDS score was associated with reduced averaged eccentricity in the somatomotor cortex and limbic network. Additionally, the DDS negatively correlated with dispersion between the DA network and somatomotor cortex, as well as between the limbic network and the somatomotor and VA networks. These correlations between a higher DDS score and more integrated limbic, somatomotor and VA networks suggest that this neural pattern is functionally relevant to understand the emotional regulatory capacities of experts. In particular, we showed on the same sample of participants that these experts were more able to reduce and to decouple the unpleasantness of a painful stimulus from its intensity than novices (10), and that the DDS was a core factor to explain the stronger sensory-affective uncoupling of pain found in experts (11).

Our study had several limitations. Its cross-sectional nature limits our ability to establish causal relationships and is biased by the self-selection bias. It is possible that some group differences presented reflect existing inter-individual differences which preceded the meditation training. The fact that cognitive defusion, a core mechanism of meditation, was associated with brain processes which overlapped with the ones related to meditation expertise, makes this interpretation unlikely. Although efforts were made to control for potential confounding variables by matching experts and novices for age, sex, and education, there may still be unaccounted factors that could explain the observed differences. In addition, our study was mainly exploratory as we used diffusion embedding to study the effect of long-term meditation practice on the brain, thus our findings will require replication by future studies.

In conclusion, we identified large-scale networks associated with meditation expertise, which were not limited to specific meditative states, and which shed new light on the neural mechanisms of cognitive defusion as measured by DDS.

## Abbreviations

OP: open presence
OM: open monitoring meditation
LKC: loving-kindness and compassion meditation
B2B: back-to-back
RS: resting-state
MBI: mindfulness-based interventions
Vis: Visual
DA: dorsal attention
VA: ventral attention
SN: salience network
FP: frontoparietal
DMN: default-mode-network
DDS: Drexel defusion scale
BDI: Beck depression inventory
FFMQ: five facets mindfulness questionnaire
DMT: N,N-Dimethyltryptamine
SM: Supplementary Materials

## Acknowledgements

We thank the expert and novice meditators for their participation in our study. The authors would like to thank the Neuropain laboratory (Lyon), Sofie Valk, Romain Quentin, Ronan Perry, and Josua T. Vogelstein for their valuable inputs in theoretical and statistical discussions. Gratitude is also extended to Clara Benson, Liliana Garcia Mondragon, Oussama Abdoun, Kristien Aarts and Eléa Perraud for their assistance during data collection, and the recruitment of experts. Appreciation is also expressed to Franck Lamberton and Camille Fauchon for their help with the technical aspects of the protocol. Their contributions have greatly enhanced the quality of this research.

## CREDIT AUTHORSHIP CONTRIBUTION STATEMENT

**Gaël Chetelat**: Supervision. **Sébastien Czajko**: Methodology, Validation, Formal analysis, Data curation, Writing, Visualization. **Loïc Daumail**: Formal analysis. **Antoine Lutz**: Methodology, Conceptualization, Supervision, Writing – editing, Funding acquisition. **Daniel Margulies**: Methodology, Supervision. **Jelle Zorn**: Conceptualization, Methodology, Investigation. Data Collection

