## Supplementaries for "Exploring the embodied mind: functional connectome fingerprinting of meditation expertise"

### Supplemental Material (SM)

### METHOD

#### PARTICIPANT:

Initial studies on expertise suggested that 10.000 hours of deliberate practice were required to reach such a stage (1). However, meta-analyses have shown that experience itself is not a sufficient criteria to quantify such a construct (2,3), as it is not always a stronger predictor than a measure related to the domain of expertise (4). Following these recommendations, we operationalized here meditation expertise by combining objective and intersubjective criteria. Firstly, practitioners had to have learned and practiced the same styles of meditations, as it is proposed in traditional 3-year meditation retreat in the Kagyu or Nyingma school of Tibetan Buddhism, where the practitioners meditate 8-12 hours a day. They also had to have the minimum requirement of 10,000 hours of formal practice as well as a regular daily practice, ensuring a certain degree of expertise of the kind of meditation practice we wished to investigate. Secondly, a research assistant (CB) highly experienced with these practices checked that the practitioners' Buddhist communities perceived them as meditators sufficiently skilled and experienced in these practices. As such, we required experts to have an extensive and regular practice in the Karma Kagyu (Mahamudra) or Nyingma (Dzogchen) schools or both. More precisely, practitioners should have accumulated $\geq$ 10000 hours of experience, followed at least one formal 3-year meditation retreat, and have a regular daily practice of $\geq$ 45 minutes in the year preceding the study. Novices should have no experience in meditation or comparable mind-body practices (e.g. yoga, tai chi, qi-gong etc.). Notable exclusion criteria regardless of group were pre-existing neurological or psychiatric conditions (e.g. epilepsy, depression) and/or any condition involving sensitization to pain (e.g. chronic pain, fibromyalgia), had no family history of epilepsy, their score to the Beck Depression Inventory (BDI) had to be below 20, had no severe hearing loss, use of medication that could interfere with relevant cognitive functions such as the central nervous system (e.g. antidepressants, opioids) or the pain system (e.g. nonsteroidal anti-inflammatory drugs). Women had not to be pregnant, breastfeeding or having given birth in the last 6 months. Subjects had to be compatible for MRI sessions, such as not being claustrophobic, not having metal implants and dental prostheses.

#### MEDITATION PRACTICE:

Mindfulness meditation consists of cultivating a vigilant awareness of one's own thoughts, actions, emotions and motivations (5). The participant learns to intentionally pay attention to his or her internal or external experiences in the present moment, without making any value judgment. The aim is that the present moment is lived in a more open and flexible way and is less dominated by mental conditioning that is a source of suffering. Two standard styles of mindfulness meditation are FA meditation and OM meditation (6). FA involves sustaining one's attentional focus on a particular object, either internal (e.g., breathing) or external (e.g., a candle flame). The practitioner is instructed to monitor their attention, notice episodes of distraction (mind-wandering), and bring their attention back to the object. OM practices aims to cultivate and sustain an effortless, open and accepting awareness of present moment experience, without changing, being reactive or absorbed in its contents. Such a dereifying perspective purportedly allows one to recognize that all components of conscious experience are simply mental events, and thus do not necessarily need to be acted upon. Thus, the aim of this training is not to explicitly change, alter or suppress experiential content, but rather to change one's relation to it. As such, an unpleasant experience might be perceived with equal or even increased vividness during an OM state, without the fear and emotional reactivity that usually accompanies such experience. "Open Presence" (OP) in Tibetan Buddhist traditions is a paradigmatic case of a so-called non-dual mindfulness meditation. Styles of meditation that cultivate OP are described as inducing a phenomenal experience where the intentional structure involving the duality between object and subject is attenuated, as captured by the notion of non-duality. OP style of practices can be found in both the Dzogchen (Tibetan, Rdzogschen) and Mahamudra or Chagchen (Tibetan, phyagchen) traditions of Tibetan Buddhist meditation [Schaik], that are highly overlapping and have as central tenet the cultivation of the OP state (Tibetan, "rig pa cog gzhag", pronounced "rigpachokshak", literally "freely resting in what consciousness manifests"). An exemplary instruction goes as follows: "Within a state free of hopes and fears, devoid of evaluation or judgment, be carefree and open. And within that state, do not linger on the past; do not invite the future; place awareness within the present, without alteration, without hopes or fears" (as quoted in (7)). Based on its traditional presentation [Namgyal, Third Dzochen Rinpoche], OP practice is considered here an advanced form of OM practice, where practitioners might have reached various stages of accomplishment. Theoretically, OP meditation consists of a state where the phenomenological qualities of effortlessness, openness, and acceptance are vividly experienced, and control-oriented elaborative processes are reduced to a minimum. Getting familiar and stabilizing the suspension of these elaborative processes requires substantial training. For this reason, the term OP will be used here only for expert meditators, even if both experts and novices received the same meditation instructions during the task. Below we use the term OM for both novices and experts, and we will assume that for experts only it will also qualify as a practice of OP. In addition, both groups practice a state of compassion meditation. Building on the non-judgmental monitoring capacity developed in OM/OP, the participants learn to cultivate kindness toward oneself for instance in relation to one's negative thoughts, distractions, difficult emotions, unpleasant physical sensations to foster appreciation toward positive qualities of one's mind (joy, contentment, …). The participants then learn to extend a similar attitude of care and loving-kindness toward their loved ones, toward neutral persons or toward difficult persons, ultimately recognizing that the need for comfort, security, and happiness is shared by all living beings. Experts referred to this practice as 'nonreferential compassion'' (dmigs med snying rje in Tibetan) or unconditional loving-kindness and compassion, which is described as an ''unrestricted readiness and availability to help living beings.' (8)

#### PARADIGM (STATES INSTRUCTIONS)

At rest, participants were instructed to keep their eyes closed and let their thoughts run freely without falling asleep. For the open-monitoring condition, participants would start the meditation by anchoring their attention in their body, while keeping both body and mind relaxed. Subsequently, they were instructed to imagine their mind as a vast and clear space and to allow any arising experiences to occur naturally without resistance, while simultaneously not engaging in any distractions. As for the compassion state, participants were asked to relax and bring to their mind the image of a close relative, and to fill their mind with love directed to this person and the wish of good for oneself, and that any suffering may be dispelled. After each fMRI scan, we asked participants to respond to self-reports of subjective experience during the different states using a 1-10 item Likert scale as follows: clarity of mind during a block, serenity -peace of mind during a block- and valence -how positive their experience was during a block.

#### PSYCHOMETRIC SCALES

The Drexel defusion scale (DDS) measures a person's ability to distance from a variety of psychological experiences using a 10-item questionnaire. The questionnaire begins with an introduction to the concept of cognitive defusion, a concept closed to dereification (9), which is intended to help respondents understand the construct. Participants were asked to indicate the extent to which they would be able to cognitively defuse themselves from hypothetical situations with negative thoughts or feelings on a 6-point Likert scale ranging from "not at all" (0) to "very much" (5). Higher scores indicate better cognitive defusion. The DDS showed good preliminary internal consistency (α = .83), and high convergent and divergent validity (10).

The Five facets mindfulness questionnaire (FFMQ) is a 39-item questionnaire that measures five purported mindfulness dimensions including: observing (noticing or attending to internal/external experiences), describing (labeling internal experiences with words), acting with awareness (attending to present moment experience), non-judging (adopting a non-evaluative stance toward thoughts and feelings), and non-reacting (allowing thoughts and feelings to pass). Participants indicate to what degree they experience these dimensions in their daily life on a 5-point Likert-type scale ranging from 1 (never or very rarely true) to 5 (very often or always true). Scores are calculated separately for subscales, with higher scores reflecting higher mindfulness. The FFMQ facets have been found to demonstrate adequate to good internal consistency, with α coefficients ranging from .75 to .91 (11).

The Beck depression inventory (BDI) is a 21-item questionnaire that measures characteristic attitudes and symptoms of depression. Each question has four scores ranging from 0 (symptom not present) to 3 (symptom very intense). A total sum score is calculated to reflect depression severity. The BDI-I has shown good internal consistency, with α coefficients of .86 and .81 for psychiatric and non-psychiatric populations, respectively, and good concurrent and discriminant validity (12).

#### Data acquisition

Neuroimaging data was collected on a 3T Siemens Prisma scanner (Erlangen, Germany) with a 64-channel head/neck coil. Functional imaging data were acquired using an EPI sequence with the following parameters: TR = 2100 ms, TE = 30 ms, number of slices = 39, slice thickness = 3.1 mm, gap between slices = 3.1 mm, voxel size = 2.8 x 2.8 x 3.1 $mm^3$, flip angle = 80°. In addition to that, the following structural images were recorded for anatomical reference: T1-weighted (TR = 2500 ms, TE = 2.83 ms, flip angle = 6˚, voxel size = 1 x 1 x 1 $mm^3$), T2-weighted (TR = 2500 ms, TE = 261 ms, flip angle = 120˚, voxel size 1 x 1 x 1 $mm^3$) and non-EPI T2*-weighted (TR = 3680 ms, TE = 30 ms, flip angle = 90˚, voxel size = 1 x 1 x 1 $mm^3$).

#### Pre-processing

Results included in this manuscript come from preprocessing performed using fMRIprep version 1.2.6-1 (13,14), a Nipype (15,16) based tool. Each T1w (T1-weighted) volume was corrected for INU (intensity non-uniformity) using N4BiasFieldCorrection v2.1.0 (17) and skull-stripped using antsBrainExtraction.sh v2.1.0 (using the OASIS template).

Brain surfaces were reconstructed using recon-all from FreeSurfer v6.0.1 (18), and the brain mask estimated previously was refined with a custom variation of the method to reconcile ANTs-derived and FreeSurfer-derived segmentations of the cortical gray-matter of Mindboggle (19). Spatial normalization to the ICBM 152 Nonlinear Asymmetrical template version 2009c (20) was performed through nonlinear registration with the antsRegistration tool of ANTs v2.1.0 (21), using brain-extracted versions of both T1w volume and template. Brain tissue segmentation of cerebrospinal fluid (CSF), white-matter (WM) and gray-matter (GM) was performed on the brain-extracted T1w using fast (FSL v5.0.9) (22).

Functional data was slice time corrected using 3dTshift from AFNI v16.2.07 (23) and motion corrected using mcflirt (FSL v5.0.9) (24). This analysis was followed by co-registration to the corresponding T1w using boundary-based registration (25) with nine degrees of freedom, using bbregister (FreeSurfer v6.0.1). Motion correcting transformations, BOLD-to-T1w transformation and T1w-to-template (MNI) warp were concatenated and applied in a single step using antsApplyTransforms (ANTs v2.1.0) using Lanczos interpolation. Physiological noise regressors were extracted applying CompCor (26). Principal components were estimated for the two CompCor variants: temporal (tCompCor) and anatomical (aCompCor). A mask to exclude signal with cortical origin was obtained by eroding the brain mask, ensuring it only contained subcortical structures. Six tCompCor components were then calculated, including only the top 5% variable voxels within that subcortical mask. For aCompCor, six components were calculated within the intersection of the subcortical mask and the union of CSF and WM masks calculated in T1w space, after their projection to the native space of each functional run. Frame-wise displacement (27) was calculated for each functional run using the implementation of Nipype. ICA-based Automatic Removal Of Motion Artifacts (AROMA) was used to generate aggressive noise regressors as well as to create a variant of data that is non-aggressively denoised (28).

Many internal operations of FMRIPREP use Nilearn (29), principally within the BOLD-processing workflow. For more details of the pipeline see https://fmriprep.readthedocs.io/en/stable/workflows.html.

#### Connectome gradient construction

Following Hong and colleagues (30), we projected grey-matter voxels on the brain surface to 10242 vertices per hemisphere and downsampled the time-series data. Then, using the fMRI time-series matrix in each subject, we calculated functional connectomes based on Pearson correlations. As in Margulies et al. (2016) (31) and other studies (30,32,33), we z-transformed and thresholded this matrix, leaving only the top 10% of weighted connections per row, and calculated a cosine similarity matrix that captures similarity in connectivity profiles between vertices. We applied diffusion map embedding (31,34), a nonlinear reduction technique, to identify principal gradient components explaining connectome variance in descending order. In this study, we followed the previous recommendation (30,31) and set $\alpha = 0.5$, a choice that retains the global relations between data points in the embedded space. We performed Procrustes rotation (https://github.com/satra/mapalign) to align components of each individual to the group-level embedding based on the Human Connectome Project S1200 sample (41).

#### Identifying co-varying factors

This method first fits a decoding model **G** on half of the dataset to find the combination of dispersion measures that maximally decodes a given trait factor. Then, it fits an encoding model **Hf** on the other half of the dataset to estimate whether the decoded **f-hat** predictions are specific to **f** and/or attributable to other covariant factors.

Interestingly, it can be shown that the coefficients of **Hf** tend to a positive value if and only if **f** is linearly and specifically coded in the measures and not reducible to its covariant factors. In other words, this means that if a trait factor is not encoded in the neural responses, there will be no reliable match between the true factor and the model prediction. Thus, it is possible to use these coefficients as a measure of specific decoding performance and test whether they are statically greater than zero or not.

In order to implement the B2B regression, we first fitted a ridge regression model **G** on a random half of the dataset, across the 35 measures of dispersion, using the RidgeCV function of scikit-learn (35). Then we again used a RidgeCV function on the remaining half of the dataset to implement the **Hf** encoding model for each of the encoded trait factor. We repeated this process 10,000 times, shuffling half the dataset at each repetition. We used the same procedure to compute a B2B regression model with permuted features, with replacement, to draw the null hypothesis distribution. Significance p-values were obtained by comparing the proportion of times the averaged coefficient of a given factor exceeds the null model coefficient.

In our case, only the DDS score had a positive value in **Hf**. To better characterize the specific dispersion measures underlying this effect, we ran 35 exploratory B2B regression analyses to predict each dispersion measure. We reported on Figure 3D, the p-value testing the hypothesis that the coefficients of **Hf** for the DDS was positive for every dispersion measure.

#### Statistical analysis

##### Voxel-wise analysis

We statistically compared gradient component scores between experts and controls using surface-based linear models implemented in SurfStat (http://www.math.mcgill.ca/keith/surfstat/) for Matlab. Surface-based findings were corrected for family-wise errors (FWE) due to multiple comparisons using a random field theory of $pFWE < 0.05$.

##### Dispersion metrics analysis

In order to show group differences through all dispersion metrics, we used the package hyppo (36) to perform a multivariate non-parametric two-sample test. To compare differences of dispersion, we computed Studentized bootstrap tests with 10,000 repetitions also called bootstrap-t test (37) rather than classical parametric Student tests. Although these tests are more computationally demanding, they are also less stringent regarding their assumptions, as they do not require assessing the normality of the sample, for example.

#### Generalization

We used the stochastic gradient descent classifier in scikit-learn (29) with a modified-huber loss to predict expertise based on the within-network dispersion, the between-networks dispersion and average dispersion of the different states or the average of the three states. The goal was to assess which state predicts the best expertise trait, as it is very likely that expertise leads to state differences, even for the RS. We used a 5-fold cross-validation to separate training and test data. We repeated this procedure 5000 times with different sets of training and test data to avoid bias for separating subjects. Because markers of expertise should be present in every state, we expected that the trained classifier trained on a given state should also be able to predict expertise if tested on the other states. Thus, we also included test batches from the other states to see how well the classifier generalized. Given the fact that we had an unbalanced number of samples, we did not use accuracy as a metric of prediction performances. We preferred the area under the curve (AUC) of the receiver operating curve to compare the prediction performances, which does not suffer from unbalanced samples. The score for each test was defined as the mean AUC of the 5000 repetitions. We tested these scores for significance by computing the null-distribution using permuted labels (38).

#### Identifying co-varying factors

In order to disentangle the respective contribution of collinear trait factors described in the Demographic (Table 1) on the measures of dispersion (Figure 2A), we computed a back-to-back (B2B) regression (39,40). This approach takes advantage of both encoding and decoding techniques: encoding models can disentangle the specific contribution of co-variable trait factors on a given dispersion measure, while decoding can combine these multiple factors into a linear model, in order to better capture the signal of interest despite a low signal-to-noise ratio. In our case, only the DDS score had a significant contribution. To better characterize the specific dispersion measures underlying this effect, we ran 35 exploratory B2B regression analyses to predict each dispersion measure. We reported on Figure 3D, the p-value testing the hypothesis that the coefficients for the DDS was positive for every dispersion measure.

10. Forman EM (2012): The Drexel defusion scale A new measure of experiential distancing. *J Context Behav Sci* 11.

May 21): Nipype: a flexible, lightweight and extensible neuroimaging data processing framework in Python. 0.13.1. Zenodo. https://doi.org/10.5281/zenodo.581704

35. Pedregosa F, Varoquaux G, Gramfort A, Michel V, Thirion B, Grisel O, *et al.* (n.d.): Scikit-learn: Machine Learning in Python. *Mach Learn PYTHON.*

36. Panda S, Palaniappan S, Xiong J, Bridgeford EW, Mehta R, Shen C, Vogelstein JT (2021, April 1): hyppo: A Multivariate Hypothesis Testing Python Package [no. arXiv:1907.02088]. arXiv. https://doi.org/10.48550/arXiv.1907.02088

37. Efron B, Tibshirani RJ (1994): *An Introduction to the Bootstrap*. CRC Press.

38. Ojala M, Garriga GC (2009): Permutation Tests for Studying Classifier Performance. *2009 Ninth IEEE International Conference on Data Mining* 908–913.

39. King J-R, Charton F, Lopez-Paz D, Oquab M (2020): Back-to-back regression: Disentangling the influence of correlated factors from multivariate observations. *NeuroImage* 220: 117028.

40. Gwilliams L, King J-R, Marantz A, Poeppel D (n.d.): Neural dynamics of phoneme sequencing in real speech jointly encode order and invariant content. 15.

41. Demirtaş M, Burt JB, Helmer M, Ji JL, Adkinson BD, Glasser MF, *et al.* (2019): Hierarchical Heterogeneity across Human Cortex Shapes Large-Scale Neural Dynamics. *Neuron* 101: 1181-1194.e13.
